# Differential effect of value and salience on saccadic reaction times

**DOI:** 10.1101/2020.03.26.009878

**Authors:** Mohammad Shams-Ahmar, Peter Thier

## Abstract

Express saccades, a mode of visually guided saccades, distinguished from regular saccades by extremely short reaction times, are triggered by inserting a temporal gap between the fixation dot and the saccade target. It is usually assumed that they are produced by a specific pathway in which the superior colliculus plays a key role. Whether and how this pathway deals with information on the subjective value of a saccade target is unknown. We, therefore, studied the influence of varying reward expectancies and compared it with the impact of the presence and absence of a temporal gap between the disappearance of the fixation dot and the appearance of the target on the visually guided saccades of two rhesus macaques (Macaca mulatta). We observed that the introduction of a gap shifted the entire saccadic reaction time distribution to shorter latencies while increasing the probability of express saccades. On the other hand, promoting the monkey’s reward expectancy shortened reaction times and increased peak velocities of regular saccades, and increased the probability of express saccades. Importantly, we observed that the reaction time and peak velocity of express saccades were not sensitive to the value of the saccade target, suggesting that the express pathway does not have access to information on value. We propose a new model on express saccades that treats the salience of visual objects in the scene differently from the subjective value assigned to them.

## INTRODUCTION

In a visually guided saccade task, inserting a temporal gap between the fixation point offset and the saccade target onset increases the probability of express saccades at the expense of regular saccades, highlighting two distinct modes of saccadic reactions (Dorris et al. 1997; Fischer et al. 1984; Fischer and Boch 1983; Fischer and Ramsperger 1984; Kingstone and Klein 1993; Mayfrank et al. 1986; Schiller et al. 2004). It is commonly assumed that these two modes arise from two—at least partially—distinct pathways, with one, responsible for express saccades, being significantly faster (Chen et al. 2013; Isa and Hall 2009; Schiller and Tehovnik 2005). The existence of these two pathways might explain the bimodality of the saccadic reaction time distribution. However, the question why such parallel pathways exist in the first place, and under which circumstances the one or the other is given preference, remains unanswered.

It is well established that both saccade reaction time and saccade peak velocity of regular saccades depend on the expected value of the saccade target (Lauwereyns et al. 2002; Milstein and Dorris 2007; Takikawa et al. 2002). Targets with large expected value are associated with shorter reaction times and higher peak velocities, features that increase the probability of getting hold of a potentially fleeing valuable target, yet, at the expense of significant investments of costly resources. On the other hand, reacting less vigorously in cases of targets of low expected value, whose loss one may bear more easily, has the advantage that these investments are lower. In any case, assessing the expected value will inevitably take time with detrimental consequences for targets that may be expected to be of existential importance. Hence, in this case, skipping the time-consuming assessment of value to react as quickly as possible – for instance to something potentially dangerous – may be the better strategy. Could it be that express saccades are a manifestation of such a strategy? Of course, to take advantage of such a selection, reliable prior assumptions on the relevance of stimuli would be needed. However, independent of the question of what their basis might be, one would expect to see no correlation between the metrics of express saccades and the expected value of the saccade target.

To test our hypothesis of a differential influence of value for express and regular saccades, we carried out a study of visually guided saccades made by well-trained rhesus monkeys. In our experiments, we manipulated saccade value and the temporal context of the presentation of the saccade target, the latter realized by randomly inserting a temporal gap between the fixation-point offset and the onset of the saccade target. Our results support the concept of two partially separated pathways, fed by common input that is responsible for a large part of the influence of low-level features of the stimuli on both. However, as hypothesized, the influence of target value is largely confined to the regular saccade pathway.

## METHODS

### Subjects and setup

Two male rhesus monkeys (8-13 kg) were trained on a task of visually guided saccades with two distinct features. First, on randomly chosen trials a temporal gap could separate fixation point offset and target onset, whereas on the other trials the two coincided. Second, monkeys received prior information on the varying amount of reward to be expected in the case of a successful trial. Eye movements were recorded with an ISCAN ETL-200 video tracker, resampled at 1 kHz. During experiments, the monkeys sat in a primate chair with their heads immobilized, 60 cm away from an LED monitor in a dim room. The surgical techniques used for the implantation of the head posts, a scleral search coil and a chamber for later recordings from the cerebellum followed protocols described in detail elsewhere (Arnstein et al. 2015). The procedure was approved by the local animal care committee under European and German law and the National Institutes of Health’s Guide for the Care and Use of Laboratory Animals.

### Behavioral procedure

Each trial began after a variable inter-trial interval of 200-500 ms, followed by a fixation period of 500 ms duration. The fixation cue, initially a tiny dot (d = 0.25°), appeared in the center of the screen and the monkey was required to keep his gaze within a window of 1.5° centering on the fixation cue. The initial fixation dot was replaced by a 1° diameter symbol for 300 ms, still requiring fixation, besides informing the monkey on the amount of reward to be expected. Two reward cues, a ring with a central dot, or a diagonal cross, indicating two possible levels of reward, where chosen at random. In both monkeys, the mapping between the two cues and the two levels of reward was reversed after about half of the experimental sessions. The reward cue lasted for another 300-600 ms. On almost half of the trials, chosen at random, its offset was followed by the immediate appearance of a target dot (d = 0.25°) at 8° eccentricity on the horizontal axis, at random left or right. On the other half of trials, the onset of the peripheral target followed the offset of the fixation cue only after a temporal gap of 200 ms during which the monkey had to keep fixating the position of the fixation dot/reward cue. Independent of the onset time of the target, the monkeys had to make a saccade to it within 500 ms following its onset and hold their gaze within the fixation window shifted to the target for 500 ms to receive a reward (Fig. 1*A*). All cues used where monochromatic had a luminance of 53 cd/m^2^ and appeared on a black background of 0.2 cd/m^2^.

**Fig. 1.**
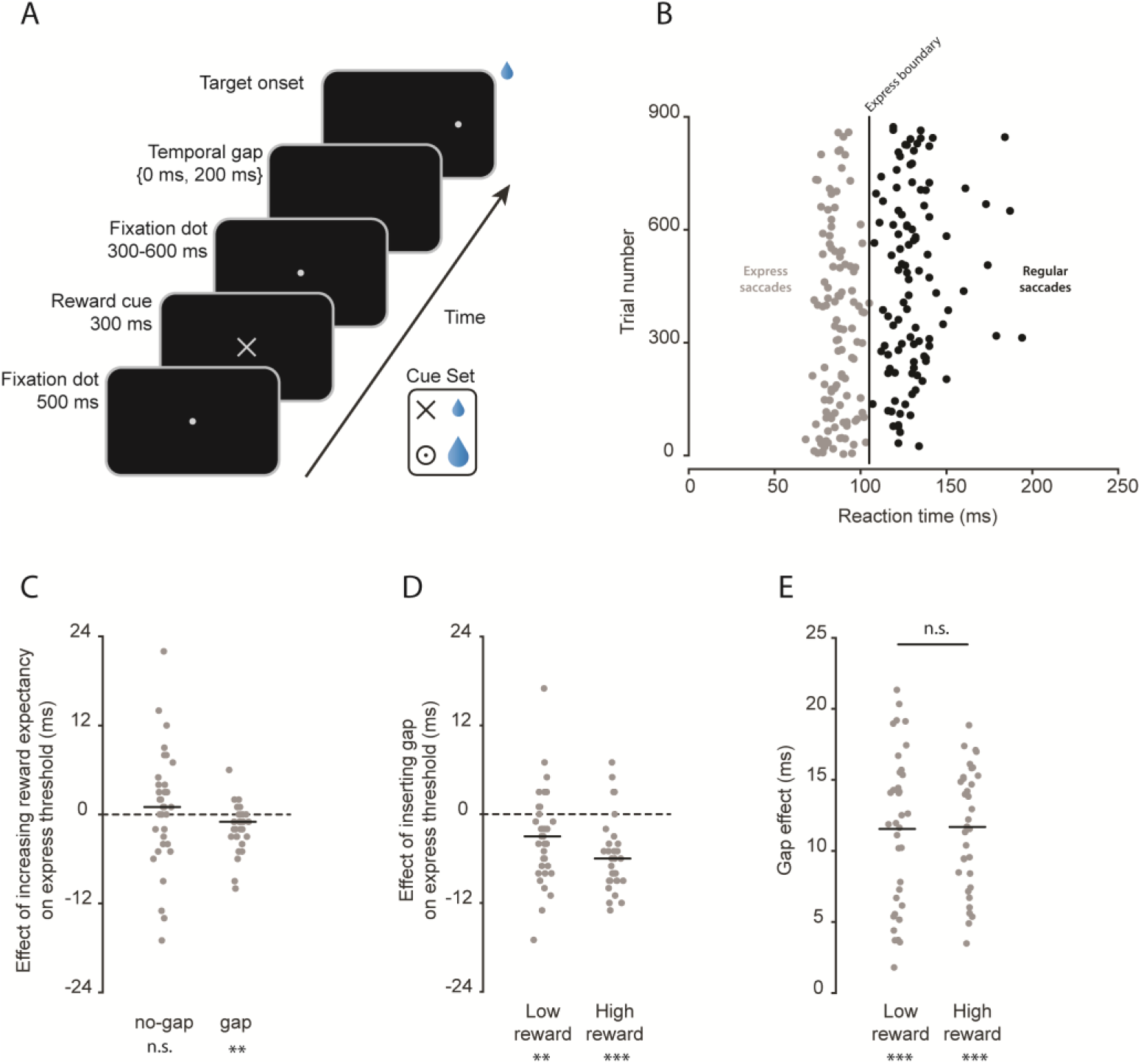
Behavioral task and the express boundary (X_bound_). *A*: the stimulus sequence. The cues were selected from the cue set and swapped their meaning after almost half of the sessions for each monkey. Dashed circles show the potential target positions. *B*: illustration of an exemplary session. The graph includes trials in gap_cond_ and LR_cond_. Each dot represents a trial and the vertical line shows the calculated X_bound_. *C*: effect of reward expectancy on X_bound_ separated by gap condition (median_gap_ = 1 ms, *p* = 0.5; median_no-gap_ = −1 ms, *p* = 0.0027). *D*: effect of introduction of a temporal gap on X_bound_ separated by reward condition (median_LR_ = −3, *p* = 0.0021; median_HR_ = −6, *p* = 10^−6^). In *C*-*D*, each dot represents an experimental session (n = 35), and the horizontal black bars indicate the median across the sessions.

To gauge the subjective value of the two reward levels possible, the standard trials were occasionally replaced by free choice trials, interspersed at random with a probability of 5%. In these free-choice trials, the high-reward and the low-reward cue appeared simultaneously on the two sides of the fixation cue, left and right axis at 8°, serving as optional targets. Their onset took place simultaneously with the offset of the central fixation dot after 500-800 ms. The monkeys could freely choose between the two target options and received the expected reward. Once, a new association of reward cues and reward level was chosen, it took the monkey 2 to 3 sessions to prefer the high-reward target in more than 90% of free-choice trials. Data from the main trials were used for the analysis of the influence of the gap and expected reward only after this choice probability had been reached.

### Eye data analysis

The eye position was recorded using the ISCAN digital eye tracker at a sampling rate of 120 Hz. To obtain a better estimation of the saccade times and positions, the eye signals were smoothed by fitting a second-degree polynomial model, using a local regression method of size 50 ms around each sample. For saccade detection, the local extrema in the velocity profile, which had absolute values between 100 to 1000 deg/s were identified as potential saccades. Then the onset and offset of each potential saccade were defined as the first data points around the extremum that fell below 25 deg/s. Saccades that landed within one degree of the targets were labeled as valid. Saccades that took less than 60 ms were considered as anticipatory saccades and were not included in further analyses.

### Distinguishing express saccades and regular saccades

To this end, we subjected saccadic reaction times to a cluster analysis using the *k*-means algorithm, with the number of clusters set to two. Reaction times in the range of 60 ms to 150 ms were fed into the clustering algorithm for potential outliers not to bias the decision of the algorithm. Clustering was run 10 times, each time with a new set of initial cluster centroids. The iteration resulting in the least sum of all the points’ distances to their corresponding centroid was taken as the final output.

### Saccade amplitude matching of peak velocities

First, the distributions of saccade amplitudes in the low-reward condition (LR_cond_) and high-reward condition (HR_cond_) were characterized by histograms with an equal bin width (0.2 ° or 0.3 °). Then, from each amplitude bin, we drew N saccades from each of the 2 distributions with N, representing the smaller of the two counts of saccades obtained for that bin for the two conditions. The corresponding peak velocities of these N trials across all amplitude bins were extracted to form the amplitude-matched distribution of peak velocities. For calculating the average peak velocity of the session, an average of 100 iterations of the same procedure was taken as the final output.

Unless otherwise stated, all statistical comparisons were based on Wilcoxon signed-rank tests (*p* < 0.001 by ***, *p* < 0.01 by **, and *p* < 0.05 indicated by *).

## RESULTS

### Differential effects of reward expectancy and the presence of a fixation gap on the express saccade boundary

We collected a total number of 35 experimental sessions (n_monkey1_ = 21, n_monkey2_ = 14). The resulting saccadic reaction time distributions of both monkeys showed distinct express saccade modes, easily discernible by eye, independent of whether a gap was present or not. Nonetheless, to avoid any subjective bias in determining the reaction time boundary between regular saccades and express saccades (X_bound_), we deployed the *k*-means clustering algorithm (see Methods). Fig. 1*B* illustrates the distribution of saccadic reaction times in an exemplary session with a gap between the disappearance of the fixation dot and the appearance of the target for low reward trials. In this particular example, the *k*-means algorithm suggested an X_bound_ separating express and regular saccades at 105 ms. The *k*-means clustering analysis was carried out separately for gap and no-gap condition saccades, furthermore separating low- and high-reward trials. Figs. 1*C*-*D* summarize the effects of the two variables reward expectancy and the presence or absence of a gap (gap_cond_ vs. no-gap_cond_) on the X_bound_: The reward level did not influence the X_bound_ in the no-gap_cond_, and reduced it only slightly, yet significantly, in the gap_cond_ (median reduction = 1 ms; Fig. 1*C*). On the other hand, the introduction of a gap caused a shift of the X_bound_ towards lower saccadic reaction times in both reward conditions. A “gap effect” i.e. the mean saccadic reaction times difference between gap_cond_ and no-gap_cond_ pooled over both saccade modes of about 12 ms was observed in both reward conditions. Thus, the introduction of a gap strongly modulated the position of the X_bound_, whereas the effect of reward expectancy was negligible.

### Differential effects of reward expectancy and the presence of a fixation gap on express saccade probabilities

We next analyzed how the reward expectancy influenced the probability of express saccades generated in the gap_cond_ vs. no-gap_cond_. In the no-gap_cond,_ both monkeys showed a significantly higher probability of generating express saccades in the HR_cond_ compared to the LR_cond_. In the gap_cond_, the same effect was obtained for one of the two monkeys and neither seen in the other monkey (*p* = 0.27) nor data pooled from both (Fig. 2*A*). One may safely assume that the subjective value of both the low and the high-level rewards attained by the monkey will steadily decrease in a session until at a point of complete satiety is reached, stopping the session. Hence, the aforementioned promotion of express saccades by large rewards might be particularly strong early in a session. In fact, this was the case. The probability of express saccades was highest in the first decade of a session and decreased steadily over sessions (Fig. 2*B*). In the presence of a gap, the probability of express saccades reached 75% in the first decade, no matter if the reward was large or small and decreased to about 50% in the last decade. In the no-gap_cond_, in which the overall probability of express saccades was less, with significantly more express saccades for larger rewards, the decrease with the trial number was much weaker, albeit still significant for the LR_cond_ and not significant for HR_cond_.

**Fig. 2.**
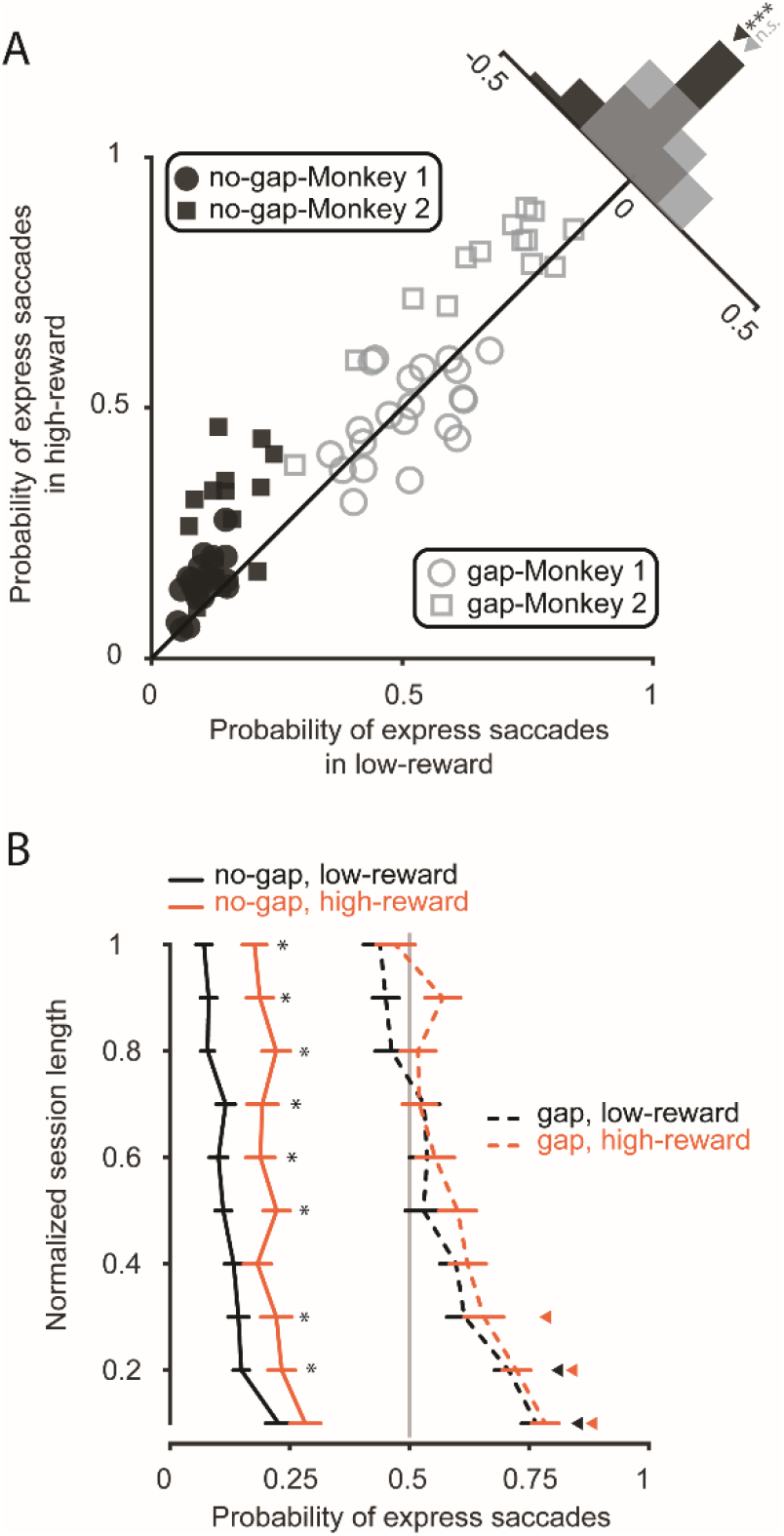
Effect of reward expectancy on the probability of express saccades *A*: comparison between the probability of express saccades in LR_cond_ and HR_cond_ in both gap conditions (median_gap_ = −0.03, *p* = 0.09; median_no-gap_ = −0.07, *p* = 2×10^−6^). Each marker represents an experimental session. The triangles above the histogram indicate the median of the corresponding distribution. *B*: the probability of express saccades throughout a session. Except for the HR_cond_-no-gap_cond_, the other trials show a significant difference of express saccade probability across trial decades (Kruskal-Wallis one-way test, LR_cond_, no-gap_cond_: *p* = 8.37×10^−7^, HR_cond_, no-gap_cond_: *p* = 0.21, LR_cond_, gap_cond_: *p* = 9.63 × 10^−13^, HR_cond_, gap_cond_: *p* = 2.03 × 10^−8^). The relative probability of express saccades in HR_cond_ was higher across trial bins in gap_cond_ (*p* = 0.0039) and no-gap_cond_ (*p* = 0.002). Horizontal lines indicate the standard error of the mean (SEM) across sessions. Asterisks indicate trial bins with significant differences between HR_cond_ and LR_cond_ (*p <* 0.05), and triangles show trial bins lying significantly above the chance level indicated by the vertical gray line (both corrected for the multiple comparison testing using Benjamini-Hochberg method).

### Differential effects of reward expectancy and the presence of a fixation gap on saccadic reaction time and saccade peak velocity

Next, we asked if manipulating reward expectancies and introducing a fixation gap influenced saccadic reaction times and the velocity of saccades. As shown in Fig. 3*A* and *C*, the saccadic reaction times of express saccades as well as their peak velocity did not show a change due to reward. However, the variability of saccadic reaction times dropped strongly (median STD difference = 1.2, *p* = 1.45 × 10^−7^). The mean of the same two factors underwent a strong modulation in the case of regular saccades, where mean saccadic reaction times dropped and mean peak velocities increased robustly due to a higher expectancy of reward. Following previous studies, we found that the peak velocity of express saccades was on average below that of regular saccades [ref]. For LR_cond_ the difference was 17.48 deg/s (*p* = 7.39 × 10^−12^) and for HR_cond_ it was 25.68 deg/s (*p* = 7.49 × 10^−11^).

**Fig. 3.**
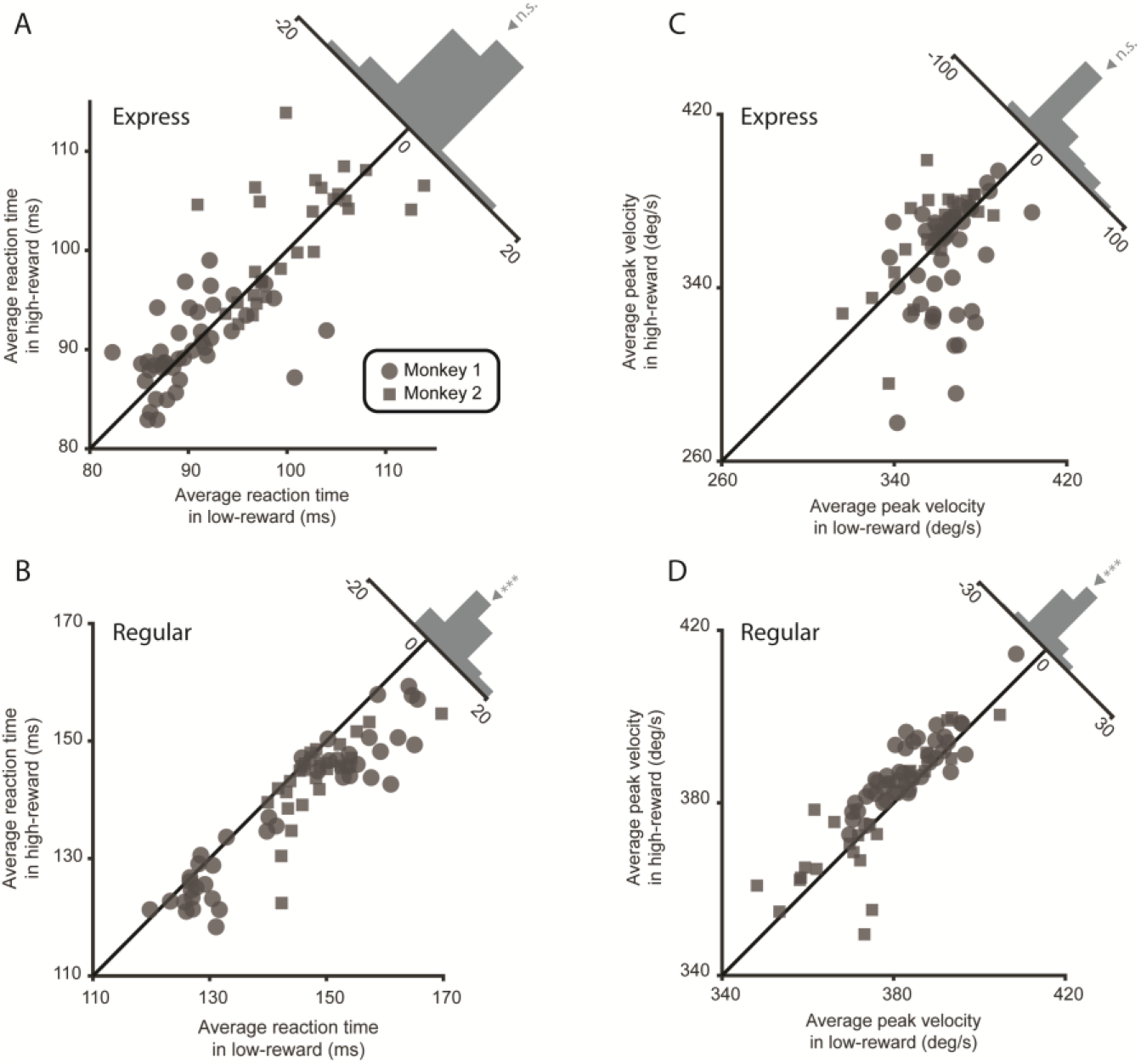
Effect of reward on saccade latency and peak velocity. *A*, *B*: comparison between the saccadic reaction time means in LR_cond_ and HR_cond_, for express saccades (*A*; median difference (HR - LR) = −1.17 ms, *p* = 0.79) and regular saccades (*B*; median difference (HR - LR) = −4.38 ms, *p* = 3.69×10^−11^). *C*, *D*: comparison between the average saccade peak velocity in two reward conditions for express saccades (*C*; median difference (HR - LR) = 2.77 deg/s, *p* = .92) and regular saccades (*D*; median difference (HR - LR) = 3.55 deg/s, *p* = 5.06 × 10^−7^). Each circle represents an experimental session. The triangles above the histograms show the median of the distribution.

### Modulation across reaction time bins

The analyses presented in the preceding paragraphs were based on the assumption of two distinct saccade modes, an express saccade mode and a regular saccade mode, and the expectation of qualitatively distinct effects of the two experimental variables reward level and the presence of absence of a gap on saccades in the two modes. The results seem to meet this expectation. However, one could argue that the a priori separation of saccades into two modes might conceal that the effects of the two experimental variables are simply graded with saccade latency, independent of a saccade interpreted as an express or regular saccade. To address this concern, we calculated the probability of saccades in individual latency bins for a pool of saccades characterized by the same reward level and the presence, or absence of a gap, bin-by-bin. Next, we subtracted the resulting probability distribution for the LR_cond_ from the one for the HR_cond_. The resulting HR_cond_-LR_cond_ distributions are shown in Fig. 4*A*, separately for the gap and the no-gap conditions. In both gap conditions, we saw that reward increased the number of saccades in the earlier saccadic reaction time bins at the expense of the later ones. In contrast to what the assumption of a graded influence of saccade latency would predict, the resulting pattern is bimodal. In the no-gap_cond_, the redistribution of later saccades to early saccades peaked at 140 ms and 100 ms, whereas in the gap_cond_ it peaked at 125 ms and 100 ms. This pattern suggests that a large reward expectancy speeded up all saccades within their respective modes while still obeying the X_bound_.

**Fig. 4.**
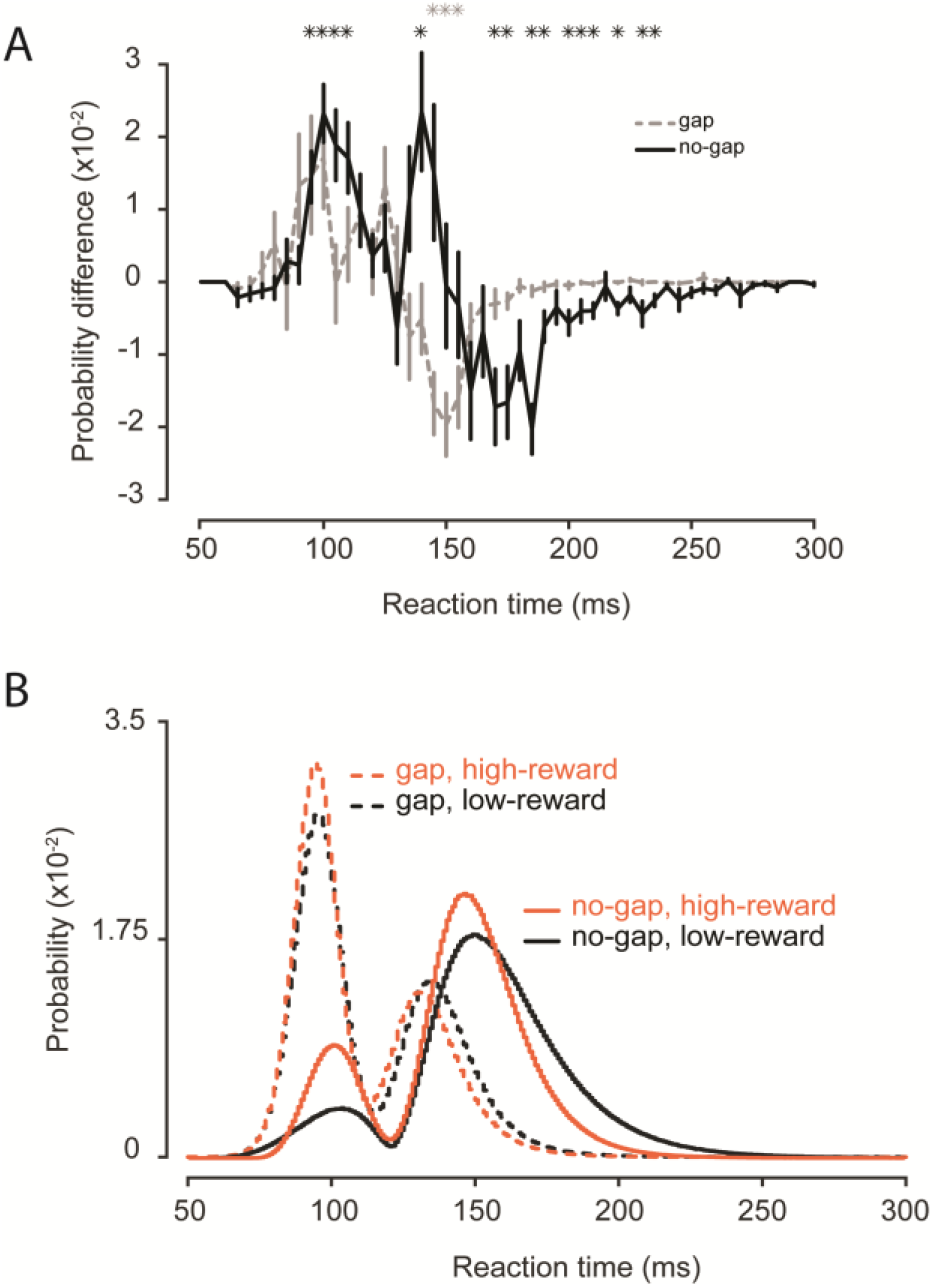
Effect of reward and gap across saccadic reaction time bins. *A*: for each gap condition, the probability of each bin in each LR_cond_ was subtracted from that of HR_cond_. Vertical lines indicate the SEM across sessions. Gray and black asterisks indicate significant bins above zero in gap and no-gap conditions respectively (corrected for the multiple comparison testing using the Benjamini-Hochberg method). *B*: the result of a simulation based on features extracted from each session using the Pearson system of probability distributions.

In order to compare the total amount of modulation between the no-gap_cond_ and the gap_cond_, we defined a modulation index for each session, given by the sum of the absolute probability differences across all saccadic reaction time bins. This comparison revealed that changes in reward expectancy had a stronger effect on saccade latencies in the no-gap_cond_ than in the gap_cond_. In summary, using a modulation index that captures both changes in express saccade proportion and in mean saccadic reaction times, implies that express saccades in no-gap_cond_ are most susceptible to the subjective value assigned to the saccade target.

Finally, to fully describe the saccadic reaction time distributions for the four conditions distinguished by reward level and the presence or absence of a fixation gap with a high temporal resolution, we quantified each resulting saccadic reaction time mode by four parameters: mean, standard deviation, skewness, and kurtosis and modeled the result by resorting to the Pearson system probability distributions. The distributions then allowed us to generate artificial trials (Fig. 4*B*), an approach that will be revisited in the discussion.

## DISCUSSION

In this study, we found different sensitivities of saccades in the express mode and in the regular mode to the expected value of the saccade. Promoting the expected value increased the peak velocities of regular saccades but not of express saccades. Furthermore, it decreased the reaction times of regular saccades up to a minimum, corresponding to the express saccade boundary. Finally, it also caused an increase in the express saccade probability. On the other hand, introducing a temporal gap shifted the express saccade boundary to shorter reaction times and increased the probability of express saccades even more.

The scheme in Fig. 5 proposes a conceptual model of express saccades that accommodates the aforementioned differential effects on regular and express saccades. The first stage of the model processes the physical properties of all visual objects, namely the fixation point, the eventual target, and the “distractors” in the visual field, independently of the subjective value assigned to them. The visual objects compete with each other to become the target of an upcoming saccade, shifting the fovea to them, based on a comparison of their “visual drive”. We assume that this comparison is based on a divisive normalization algorithm (Carandini and Heeger 2012; Louie and Glimcher 2019). The visual drive or salience of individual visual objects is obtained by dividing the individual salience by the sum of the saliencies of all objects, resulting in values between zero and one. In a natural setting, the object associated with the largest salience will win and correspondingly bind the observer’s foveal attention. The larger the salience of the winning object is, the shorter the saccadic reaction time will be. In a scenario in which just two objects staggered in time are seen, the first one available will bind the fovea first, i.e. serve as fixation dot, while the latter will serve as the saccade target. The advanced removal of the first one, the fixation dot in our paradigm, will maximize the salience of the later object, the saccade target.

**Fig. 5.**
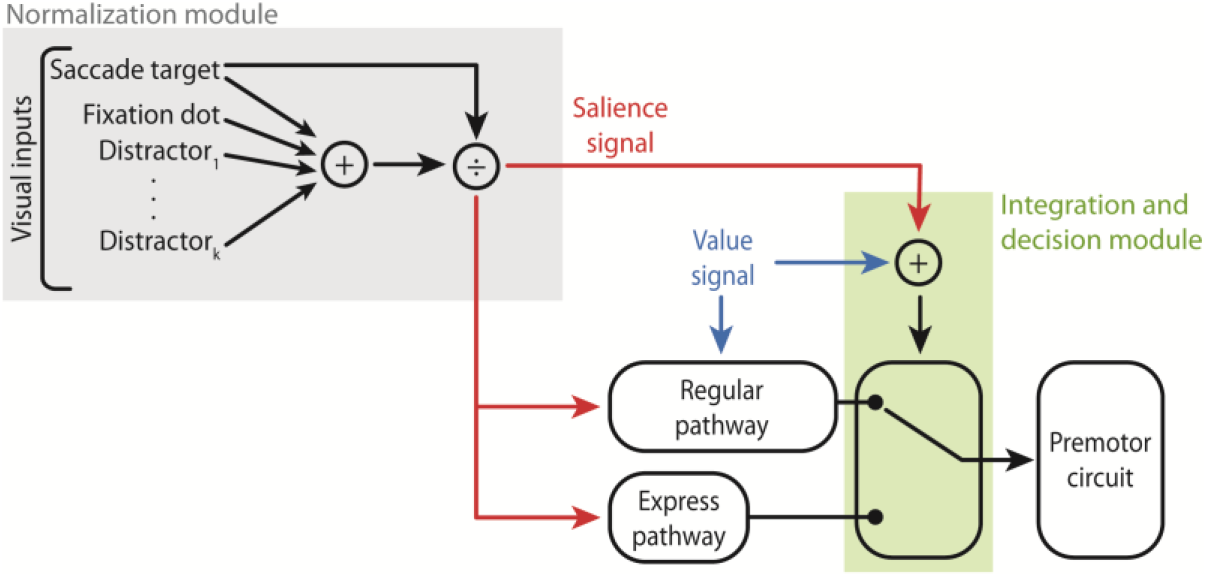
Partially separated pathways for express and regular saccades. The physical properties of the saccade target are assessed by a divisive normalization process (gray box) that results in a salience signal of the saccade target (red arrows). This signal reaches the express saccade pathway and as well the regular saccade pathway. The latter involves a receiver of value signals (blue arrows). An integration and decision module (green box), made of a switch under the joint influence of salience and value signals, decides which of the two pathways gets access to the premotor circuitry for saccades.

The salience signal feeds two parallel pathways, namely the regular and the express saccade pathway, thereby influencing all saccades, i.e. shaping the entire reaction time distribution, based on express and regular saccades. Information processing based on the regular pathway takes longer than the processing accommodated by the express pathway. The reason is that the former has to incorporate information on the value to adjust the timing and the peak velocity of regular saccades according to the subjective value assigned to the saccade target. The express pathway is faster because it skips this process, remaining indifferent to the value of the saccade target. Hence, value affects the regular saccade mode of the reaction time distribution, while not touching the express saccade mode. Whether information is transferred to the premotor circuits from the one or the other pathway is determined by a switch that is controlled by the joint influence of salience and value signals. As a result, both value and salience signals influence the probability of express saccades.

Several pieces of evidence suggest that the intermediate layers of the superior colliculus (SCi) might be a good candidate to serve as the integration and decision module in the scheme (Fig. 5, the green box). First, they receive visual information from the retina and visual cortex via superficial layers of the superior colliculus (Coe and Munoz 2017). Hence, the salience information might originate from these structures. Alternatively, it might be derived from posterior parietal cortex, which has direct projections to SCi (White et al. 2017; Wolf et al. 2015). Second, the basal ganglia, a system that plays a central role in encoding value, communicates with the SCi via the substantia nigra pars reticulata (Ikeda and Hikosaka 2003; Munoz and Everling 2004). And third, unidirectional ablation of SC eliminates express saccades in the contralateral direction (Schiller et al. 1987). Other studies have shown that the amount of build-up activity of visuomotor neurons in the deeper layers of the superior colliculus during the gap is a good predictor of the occurrence of an express saccade (Dorris et al. 1997). This build-up activity is controlled by target predictability (Basso and Wurtz 1997, 1998), as well as by target brightness (Marino et al. 2015). Whether value signals are represented in the SCi or deeper layers during the gap, and the question of how the integration of salience and value takes place in those layers, will require further experimental and computational studies.

## ACKNOWLEDGMENTS

The authors would like to thank Dr. F. Bunjes for implementing the behavioral paradigm, and Dr. P. Dicke, for assisting in the surgical procedure.

